# Shared and distinct cortical mechanisms for working memory and decision-making

**DOI:** 10.1101/2025.05.21.655340

**Authors:** Shanna K. Murray, Daeyeol Lee, Hyojung Seo

**Affiliations:** Department of Psychiatry, Interdepartmental Neuroscience Program, Yale School of Medicine, New Haven, CT 06510; The Zanvyl Krieger Mind/Brain Institute, Department of Neuroscience, Psychological and Brain Sciences. Kavli Neuroscience Discovery Institute, Johns Hopkins University, Baltimore, MD, 21218; Department of Psychiatry and Neuroscience, Yale School of Medicine, New Haven, CT 06510

**Keywords:** Prefrontal Cortex, Posterior Parietal Cortex, Reinforcement Learning

## Abstract

The dorsolateral prefrontal cortex (DLPFC) and lateral intraparietal cortex (LIP) in the primate brain are critically involved in working memory during tasks that require the retention of information over a delay. These same regions have also been implicated in reinforcement learning (RL), where information about an animal’s choice and its outcome is retained to update future reward expectations based on past experiences. We investigated whether spatial memory, required across different behavioral contexts, relies on a shared neural mechanism. To explore this, we analyzed neural activity recorded from rhesus monkeys engaged in three distinct tasks—the oculomotor delayed response task (ODR), a visual search task, and the matching pennies game—each requiring the retention and use of similar spatial information under different cognitive demands. The ODR task demands only prospective memory, as the selection of action is dictated by the location of visual cue, and the subject must retain this intended action for execution after a temporal delay. In contrast, the matching pennies task engages both retrospective and prospective memory: retrospective memory of previous choice and its outcome to inform decision-making, while prospective memory is needed to carry out that decision. Visual search task, by comparison, does not explicitly require either retrospective or prospective memory.

Our analysis revealed that neural signals encoding retrospective memory of the animal’s choice in the visual search and matching pennies tasks were not correlated with the prospective working memory signals of visually cued locations in the ODR task, in either the DLPFC or LIP. Moreover, retrospective choice signals in the visual search and matching pennies tasks were not correlated with each other. In contrast, neural activity related to upcoming choices (prospective memory) in the LIP showed significant correlations across all three tasks. In the DLPFC, prospective choice signals were correlated between the visual search and ODR tasks, but not between those tasks and matching pennies. Additionally, in the DLPFC, neural signals representing previously rewarded choices were significantly correlated with working memory signals during the ODR task.

These results suggest that the LIP supports a consistent, shared mechanism for prospective memory linking a committed action to its eventual execution. In contrast, the DLPFC might mediate the transformation of retrospective memory-integrating past choices and outcomes – into a decision and its associated prospective memory.

## Introduction

Learning about the outcomes expected from alternative actions is a critical skill required for decision-making by humans and other animals. In neuroscience, reinforcement learning theory has provided a critical framework for modeling decision-making behavior and studying its neural correlates (Sutton & Barto, 1998). Work in this field has shown that neurons in numerous brain regions such as prefrontal cortex, parietal cortex, and striatum represent task variables necessary for reinforcement learning (Schultz et al., 1997; Barraclough et al., 2004; O’Doherty et al., 2003; Samejima et al., 2005; Kim et al., 2009; reviewed in Lee et al., 2012). In particular, single neurons in regions such as dorsolateral prefrontal cortex (DLPFC), supplementary eye field (SEF)/dorsomedial prefrontal cortex (DMPFC), and lateral intraparietal cortex (LIP) encode mnemonic representations of previous choices over multiple trials (Seo et al., 2007; Seo et al., 2009; Seo & Lee 2009; Donahue et al., 2013). In the same regions, neurons represent memories of previous rewards or outcomes, and neural activity may be modulated by both choices and outcomes independently as well as their non-linear interaction.

In the field of working memory, it is well established that cortical neurons display directionally-tuned, persistent activity while items are held in memory for up to a few seconds. These neurons have been demonstrated to be necessary for working memory in physiology studies (Goldman-Rakic 1995; Funahashi et al., 1989; Fuster & Alexander, 1971). Neurons with persistent delay activity are found in multiple cortical regions including DLPFC and LIP. Evidence for persistent activity in DLPFC has been observed in human fMRI experiments as well (Curtis & Esposito, 2003). More recently, studies have suggested that persistent activity of subpopulations of single neurons is but one aspect of a more dynamic population-level representation (Cavanagh et al., 2018; Murray et al., 2017). This representation may vary depending on task constraints, allowing the brain to support stable but flexible working memory.

It is not well understood how the same neural populations support multiple cognitive functions such as working memory and reinforcement learning. However, a number of studies using recurrent neural networks have begun to build an understanding of the computational requirements of a system which can flexibly solve multiple tasks (Johnston & Fusi, 2023; Ehrlich & Murray, 2022; Yang et al., 2019; Wang, 2008). This work has highlighted the important role of non-linear mixed-selective neurons in these flexible networks, since neurons that multiplex multiple task-relevant representations within and across tasks lead to higher population dimensionality, enabling more linear readouts from the network (Rigotti et al., 2010). These findings *in silico* are consistent with work in physiology which has demonstrated the importance of neurons with mixed selectivity in solving tasks that require flexible yet stable representations (Rigotti et al., 2013; Parthasarathy et al., 2017). The prefrontal cortex is particularly rich with neurons which display mixed-selective properties, which may allow it to carry out its diverse functions related to flexible behavior and executive function. More recently, there is also evidence that mixed selectivity is a signature of participation in highly abstract representations which are apparent only when their activity is examined across multiple tasks (Ehrlich & Murray, 2022).

A small number of human studies have begun to specifically investigate the relationship between working memory and reinforcement learning (Collins, 2018; Collins & Frank, 2012; reviewed in Curtis & Lee, 2010). These studies have focused on working memory and reinforcement learning as fully separable constructs. However, the extent to which the same neurons display analogous responses across behavioral tasks is not well-studied. Single neurons are typically recorded during a single task, and even when they are tested for multiple tasks additional tasks usually serve as controls to pinpoint a particular cognitive construct. There are some exceptions. For example, some studies of the DLPFC have explored the extent to which neural activity during delays is related to visual attention or to memory per se (Lebedev et al., 2004; Messinger et al., 2009). One study found evidence that delay activity in the DLPFC represents a domain-general mnemonic signal (Tsujimoto et al., 2012). Furthermore, it is typically assumed that the same neurons will display comparable visuomotor properties across tasks, although this is an open empirical question.

Single neurons involved in maintaining memories of previous actions and action-outcome conjunctions for reinforcement learning and decision-making are found in the same regions as those supporting working memory (Lee et al., 2012). Therefore, we asked whether the same neurons may encode relevant information across working memory and reinforcement learning in a similar way. Specifically, we hypothesized that individual neurons in DLPFC and LIP would congruently encode information for working memory and reinforcement learning. To test this hypothesis, we analyzed a previously collected dataset in which the same single neurons were recorded in two decision-making tasks, each with different behavioral requirements, and one working memory task. Comparisons of activity between these three tasks have not previously been performed or published. We compared the neural responses between the working memory task and each decision-making task. While basic visuomotor properties were largely consistent for the same neurons across decision-making and working memory tasks, the way single neurons signal retrospective and prospective memory of animal’s choice varied significantly depending on behavioral contexts, and between DLPFC and LIP.

## Methods

### Animal preparation

Three rhesus monkeys weighing between 5 and 11 kg (male: H and I; female: K) were trained on the three different behavioral tasks described below. The animals sat in primate chairs and were head-fixed during the experiment. Eye movements were sampled at 225 Hz using a high-speed eye tracker system (Thomas Recording). All procedures in this study were approved by the University of Rochester Committee on Animal Research and adhered to the Public Health Service Policy on Humane Care and Use of Laboratory Animals and the *Guide for the Care and Use of Laboratory Animals*.

### Behavioral tasks

The dataset analyzed in this study was obtained using three different oculomotor behavioral tasks shown in **Figure 1**. The oculomotor delayed response (ODR) task provided information about each neuron’s directional tuning properties for working memory and for eye movements directed towards the remembered locations (**Fig. 1C**). At the beginning of each trial, the animal fixated on a centrally presented yellow square (size 0.9° x 0.9°) for a 0.5s fore-period. During the 0.5-s cue period, a circular green target (radius 0.6°) was presented at one of eight different peripheral locations 5° away from the central fixation square while the animal maintained central fixation. The target then disappeared, and the animal was required to maintain central fixation during a 1-s memory period. When the central fixation square disappeared at the end of the memory period, the animal made a saccade to the peripheral region (radius 3°) where the target was originally presented. Correct responses were rewarded with a drop of juice.

**Figure 1.**
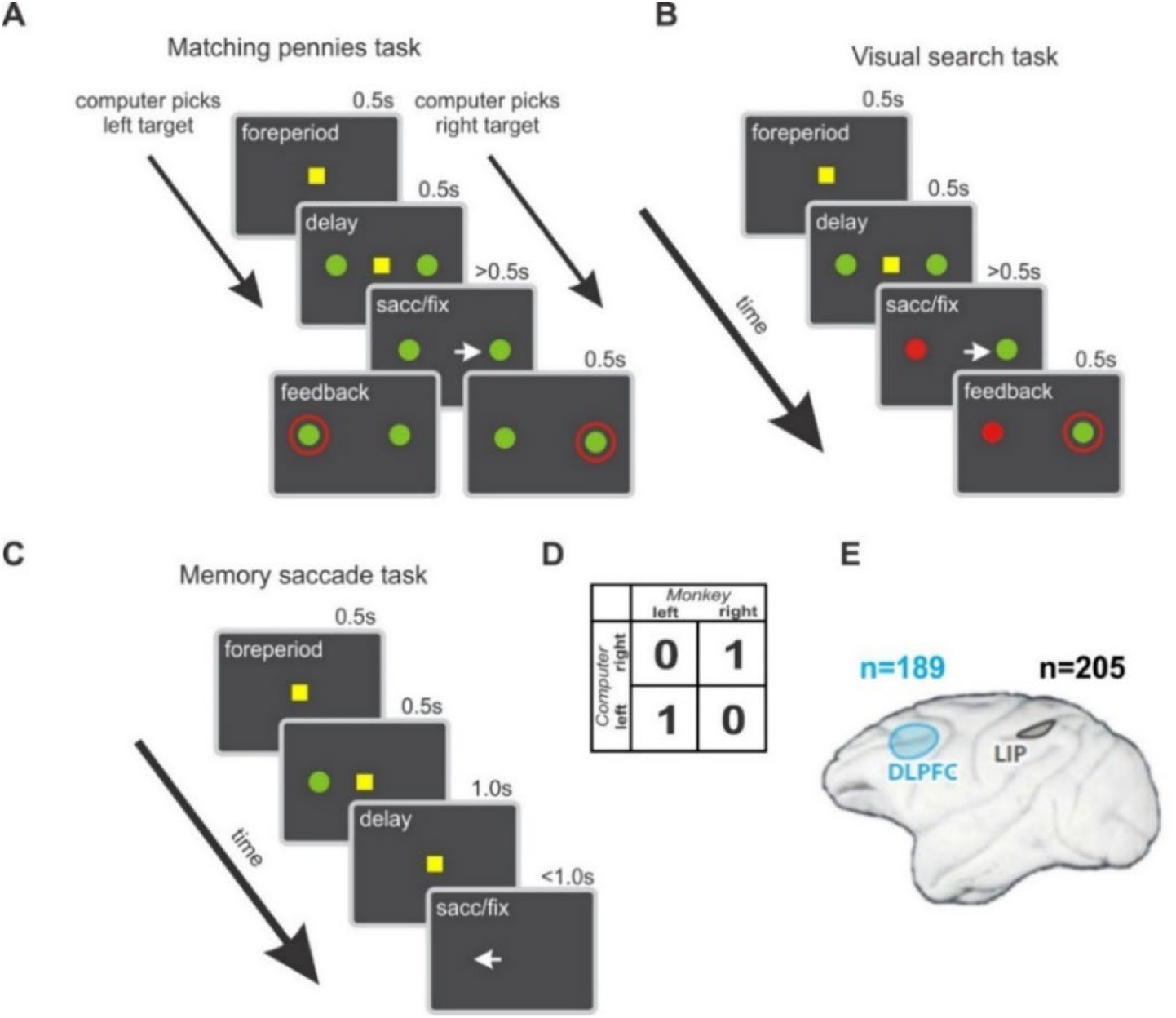
Behavioral tasks and neurophysiological recording regions. **A.** Matching pennies task. Animals played a competitive decision-making game against a computer algorithm opponent. **B.** Visual search task. Animals learned to select the green target (*p*_reward_ = 0.5) on each trial and avoided the unrewarded red target **C.** ODR task. Animals were required to remember a cued location during a 1s delay period and report the cued location at the end of the delay. **D.** Matching pennies payoff matrix. Animals were rewarded when they picked the same target as the computer. **E.** Neurophysiological recording. Well-isolated single units were recorded during all three tasks (DLPFC: n = 189 neurons; LIP: n = 205 neurons).

The matching pennies task was a zero-sum game in which the animal was rewarded for picking the same target as a simulated computer opponent (**Fig. 1A**). The computer algorithm was modeled after a rational decision-maker and tracked the animal’s choice and reward history from the session. If the algorithm did not detect any significant biases in the animal’s behavior, it chose randomly, and so the optimal strategy for the animal was to pick randomly. If the algorithm detected systematic statistical biases in the animal’s choice behavior, it exploited these biases to minimize the animal’s payoff. For example, if the algorithm detected that the animal tended to switch from left to right after not being rewarded, the computer would increase its chance of staying on the left target on trials where the animal previously chose left and was not rewarded. As in the ODR task, the matching pennies task also began with a 0.5s fore-period where the animal fixated on a central yellow square (size 0.9° x 0.9°). Two circular green targets (radius 0.6°) then appeared 5° to the left and right of the central fixation square while the animal continued to fixate for 0.5s. At the end of this delay period, the fixation square was extinguished, and the animal made a saccade to either the left or right target to indicate its choice within 1s. The animal was required to maintain fixation on the chosen target for 0.5s. The animal then received visual feedback in the form of a red ring (radius 1°) that appeared for 0.5s around the target chosen by the computer opponent. The animal received a juice reward only if it chose the same target as the computer opponent.

The visual search task events and timing were identical to the matching pennies task (**Fig. 1B**). The stimuli were identical with the exception that one of the two circular targets was red instead of green. The position of the two colors was pseudorandomized on each trial so that each target had an equal probability of appearing at the left or right location. The position and outcome for the green target were balanced so that each of the 64 possible combinations of target location and reward in a sequence of three trials was presented twice, with two leading trials to account for the lags analyzed (130 total trials). The red feedback ring appeared around the green target during the feedback period of each trial, and the animal received a juice reward with 50% probability if it chose the green target.

### Neurophysiological recording

The activity of single neurons was recorded in two different cortical regions (DLPFC, and LIP) using a five-channel multielectrode recording system (Thomas Recording). All DLPFC neurons were recorded anterior to the frontal eye field. The position of the frontal eye field was determined in each animal using <50 μA electrical stimulation to evoke eye movements. LIP neurons were recorded along the bank of the intraparietal sulcus at a minimum depth of 2.5mm from the cortical surface. Neurons were included in the analysis when they were isolated for 80 trials (10 trials per spatial location) in the ODR task, 130 trials in the visual search task (2 trials each for 64 conditions plus two leading trials), and a minimum of 130 trials in the matching pennies task.

### Behavioral data analysis

#### Logistic regression

Behavior from the two decision-making tasks was analyzed with the following logistic regression model predicting the choice location (left versus right) on the current trial *t*:

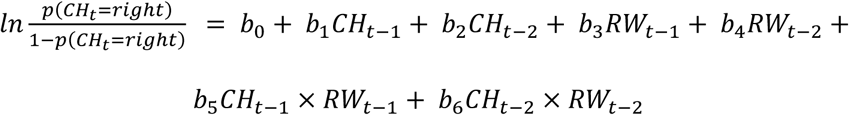

where *CH* indicated the choice at a given subscripted trial lag coded as 1 and 0 for right and left, respectively, *RW* indicated an outcome at given lag coded as 1 and 0 for rewarded and unrewarded, respectively. The interaction between a previous choice and outcome corresponded to the previously rewarded choice, which in the matching pennies task was equivalent to the computer’s choice. Results for longer trial lags have been previously published (Barraclough et al., 2004; Seo & Lee, 2007; Seo et al., 2007; Seo et al., 2009), and showed that sequential choices and outcomes in the matching pennies task were only weakly correlated. We applied the same model to the behaviors during the visual search task only to confirm that the choice and reward history was properly balanced. Significance of regression coefficients was determined with a t-test (two-tailed) on the regression coefficients with α = 0.05. Coefficients for previous choices and previous choice-outcome conjunctions were compared between the two decision-making tasks using two sample t-tests with a corrected α = 0.05/4 = 0.0125 (two-tailed) which accounted for comparisons of the two variables across the two tasks.

### Neural data analysis

#### Multiple regression

Encoding of different task-related variables for each task and neuron were quantified using multiple regression. Standardized regression coefficients were calculated for each neuron and task using z-scored regressors and z-scored spike data.

For the ODR task, this took the form of a circular regression model which contained regressors for the horizontal and vertical components of the spatial position of the remembered target in degrees as well as a constant term. Spike counts for each trial were determined for each epoch of interest and modeled according to the following equation:

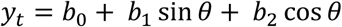

For the correlations between tasks, the 1-s time window starting 100 ms from the delay period onset was used as the epoch of interest in order to minimize the impact of stimulus offset on the estimates of working memory direction selectivity. The coefficient representing the extent to which the neuron’s activity was modulated by the horizontal position of the target held in memory (*b*_2_) was used in task-task correlations.

Two identical models were used to quantify task variable encoding for matching pennies and visual search. These models included regressors for current choice and previous choices, current and previous outcomes, and the current and previously rewarded choices (interactions between choice and reward). Spike counts for each trial were totaled for each epoch of interest examined and modeled according to the following equation:

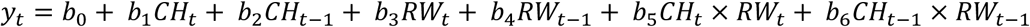

where subscripts t and t-1 index specific trials, CH represents choices coded as -1 and 1 for left and right, respectively, and RW represents outcomes coded as -1 and 1 for unrewarded and rewarded, respectively. For the initial task-task correlations, the delay period of the task was used as the epoch of interest.

To examine the correlation between regression coefficients for the two decision making tasks (referred to as task-task correlation), standardized regression coefficients were compared and analyzed across tasks using Pearson correlations corrected for multiple comparisons, for which three behavioral variables (previous choice, previously rewarded choice, and upcoming choice), two decision-making tasks, and two brain regions were considered, yielding α = 0.05/12 = 0.004 (two-tailed). Significant encoding of task-related variables was determined with t-tests of standardized regression coefficients with α = 0.05 (two-tailed). Fisher’s z-tests were used to compare correlation coefficients *ad hoc* and therefore used α = 0.05 (one-tailed).

#### Visual-motor index calculation

A visual-motor index (VMI) based on Ray et al. (2009), quantifying the extent to which a neuron’s firing rate was differentially affected by visual stimuli and saccade preparation in each task, was quantified as follows:

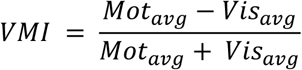

where *Mot_avg_* was defined as the average number of spikes in the 100ms before to 50ms after the beginning of the response period across all trials in a given task and *Vis_avg_* was defined as the average number of spikes in the period 50 to 200ms after stimulus onset for all trials in a given task. If there were no spikes in either period, the VMI was recorded as 0 for that neuron and task. Positive VMIs indicated that a given neuron displayed more movement-related activity than visually-evoked activity.

Comparisons between VMIs across working memory and decision-making tasks used Pearson correlations with α = 0.05 / 4 = 0.012, correcting for comparisons of the two decision-making tasks for two regions. Fisher’s z-tests were used to compare correlation coefficients *ad hoc*, using α = 0.05 (one-tailed).

## Results

The electrophysiological activity of 394 well-isolated single neurons was recorded from three rhesus monkeys (H, I, K) during three different behavioral tasks (DLPFC: 189 cells LIP: 205 cells; **Fig. 1E**). In the ODR task (**Fig. 1C**), the animals were shown a cue for 0.5s at one of eight possible peripheral spatial locations. After the cue period, they were required to remember the cue during a 1s delay period, after which they reported the location of the cue with a saccade to the remembered location.

In the two decision-making tasks, the animals were shown two visual stimuli displayed directly to the left and right of the center during a 0.5s delay period. After the delay period, they reported their choice with a saccade to and fixation on one of the two stimuli. Feedback was then displayed on the screen in the form of a red ring around the rewarded target. In the visual search task (**Fig. 1B**), one stimulus was green and the other red. The spatial positions of the green and red target during the visual search task were pseudo-randomized across trials. Critically, only the green target was rewarded (*p*=0.5), and the red target was never rewarded. Accordingly, the animals did not need to remember the most recent choice locations or resulting outcomes in order to make decisions in the visual search task. By contrast, the matching pennies task (**Fig. 1A**) was a highly dynamic decision-making task in which the animals played a zero-sum game (**Fig. 1D**) with a simulated computer opponent. The computer algorithm tracked the full choice and reward history from the task session. If the algorithm detected any bias in the animals’ choice behavior, it exploited this bias to reduce the animal’s reward rate. If no bias was detected, the computer chose randomly. The optimal strategy for the animals was therefore to choose randomly.

Nevertheless, as described previously (Barraclough et al., 2004; Seo & Lee, 2007; Seo et al., 2007; Seo et al., 2009; Donahue et al., 2013), the animals’ choices in matching pennies are weakly but systematically affected by the recent history of choices, outcomes, and their conjunctions, consistent with reinforcement learning. These matching pennies behavioral results for the three animals in the present dataset are replicated in **Figure 2, A-B** (solid lines). For the visual search task, as expected, choices were not affected by the recent choice locations or by recent choice-reward conjunctions (**Fig. 2. A-B**, dashed lines). All four sets of coefficients were significantly larger in magnitude for matching pennies compared to visual search (two-sample t-tests on regression coefficients; all *p* < 5.0e-13, two-tailed). Although information about previous choices and choice-outcome conjunctions was used for decision-making in the matching pennies task only, we found that a significant proportion of neurons in both DLPFC and LIP encoded information about the current and previous choice location, current and previous reward and the two most recently rewarded choices in both tasks (**Fig. 2. C-D**). To test our hypothesis that information may be encoded in a similar way between decision-making and reinforcement learning, we focused on the relationship between the mnemonic representations from the ODR task and each of the decision-making tasks.

**Figure 2.**
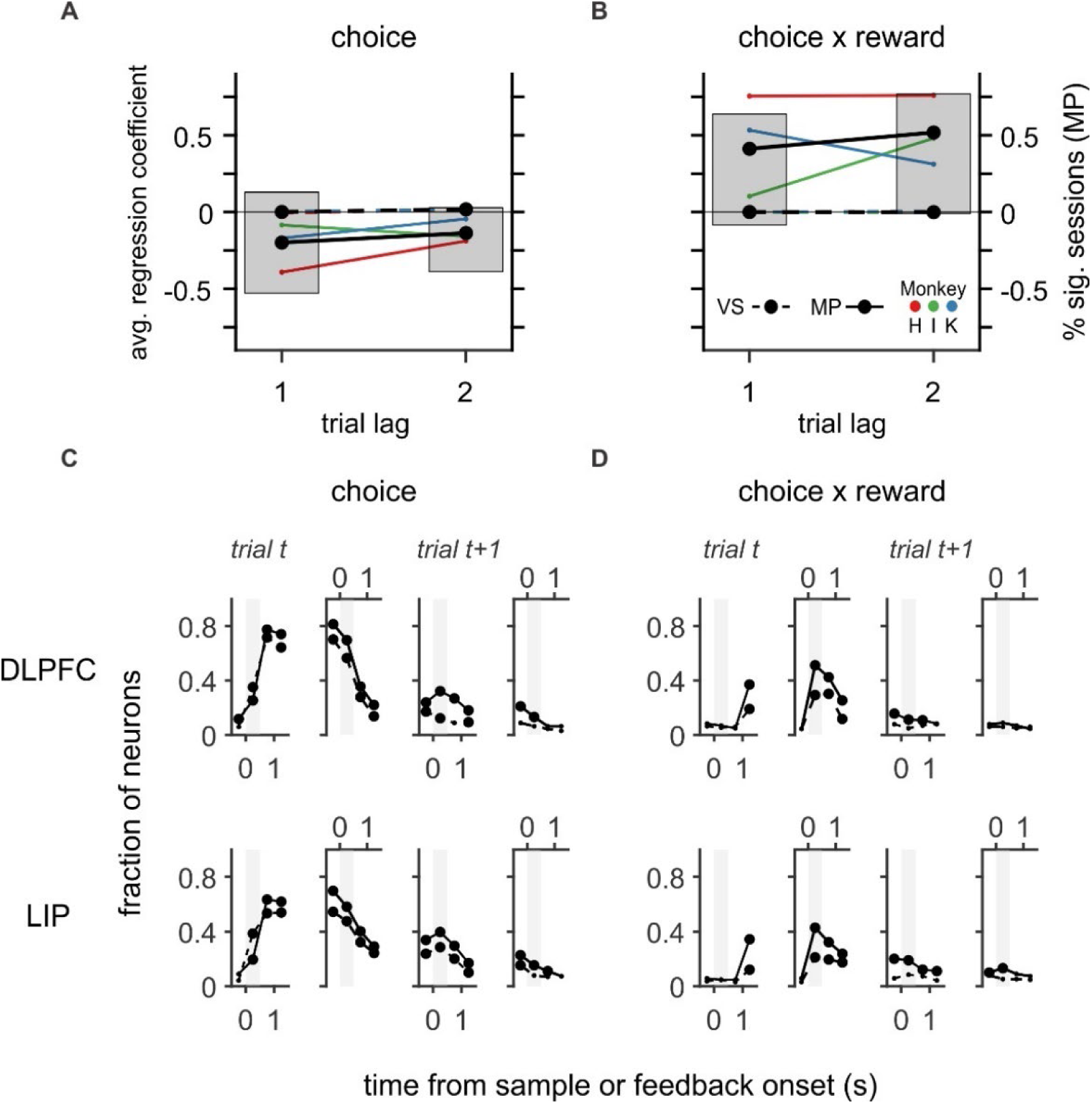
Behavioral results and neural encoding of choice variables in DLPFC and LIP during decision-making. **A-B.** Regression coefficients from the matching pennies (MP; solid lines) and visual search tasks (VS; dashed lines) showing the effect of previous choices (A) and previous choice x reward interactions (B) on decision-making as well as the fraction of MP sessions with significant coefficients. Coefficients were not significant for any variable or session for VS. **C-D**. Fraction of neurons in DLPFC (top) and LIP (bottom) encoding the choice from trial *t* in (C) or the choice x reward interaction (rewarded choice) from trial *t* in (D) on trial *t* and on the subsequent trial *t+1*. Encoding was estimated with multiple regression and is aligned to sample and feedback on each trial. Larger markers indicate the fraction of neurons was significantly larger than expected by chance (0.05; one-tailed binomial tests corrected for multiple comparisons). Dotted and solid lines represent visual search and matching pennies task, respectively.

We first quantified the tuning of neurons in DLPFC and LIP to the previous choice and previously rewarded location in the two decision-making tasks using standardized multiple regression. Encoding of these memory representations in these tasks was quantified for the delay period during which the animals made their decisions. To obtain a comparable measure for working memory, we quantified the tuning of single neurons in the two regions along the horizontal axis during the delay period of the ODR task. Because neurons in these regions respond to visual input with a brief delay, both epochs were shifted by 100ms to avoid activity related to changes in visual input at the beginning of the epoch (decision tasks: 100ms to 600ms after stimulus onset; ODR task: 100ms to 1100ms after stimulus offset). Importantly, we found that overall firing rates in these epochs of interest were not significantly different between the working memory task and decision-making tasks in either region (data not shown; DLPFC: ODR-matching pennies (MP), *p* = 0.15; ODR-visual search (VS), *p* = 0.089; LIP: ODR-MP, *p* = 0.47;ODR-VS, *p* = 0.43; t-tests, two tailed), and overall firing rates in these epochs were highly correlated between the working memory and decision-making tasks (data not shown; all Pearson’s *r* > 0.80, *p* < 6.9e-44; two-tailed). Regression models for all tasks were designed such that negative standardized regression coefficients (SRCs) indicated leftward tuning and positive SRCs indicated rightward tuning. Using these measures, we asked whether at the level of the population, neurons consistently encoded items in working memory and remembered information for decision-making.

Given that many neurons in both regions represent the previous choice during the delay period of the decision-making tasks, we first asked whether there was a significant correlation between SRCs representing the previous choice in the decision-making tasks and SRCs for horizontal mnemonic tuning for the tasks in DLPFC and LIP. We found that there was no significant relationship between the SRCs for working memory and either decision-making task in DLPFC or LIP (**Fig. 3A-D**; all *p* > 0.1, two-tailed). We next asked whether there was a significant correlation between SRCs representing the previously rewarded choice (equivalent to interaction between choice and reward) in the decision-making tasks and the SRCs for horizontal mnemonic tuning in the two regions. The previously rewarded choice was highly relevant for decision-making in the matching pennies task, but not in the visual search task. We found that SRCs for the previously rewarded choice in matching pennies and SRCs for horizontal working memory encoding were significantly correlated in DLPFC (**Fig. 4B**; Pearson’s *r* = 0.25*, *p* = 4.4e-4; two-tailed; corrected for multiple comparisons). No significant correlation was observed for the visual search task (**Fig. 4D**; Pearson’s *r* = 0.13, *p* = 0.087, two-tailed). However, the correlation between SRCs for working memory horizontal tuning and SRCS for the previously rewarded choice was not significantly larger for matching pennies compared to visual search (Fisher’s *z*-test; *p* = 0.11, one-tailed).

**Figure 3.**
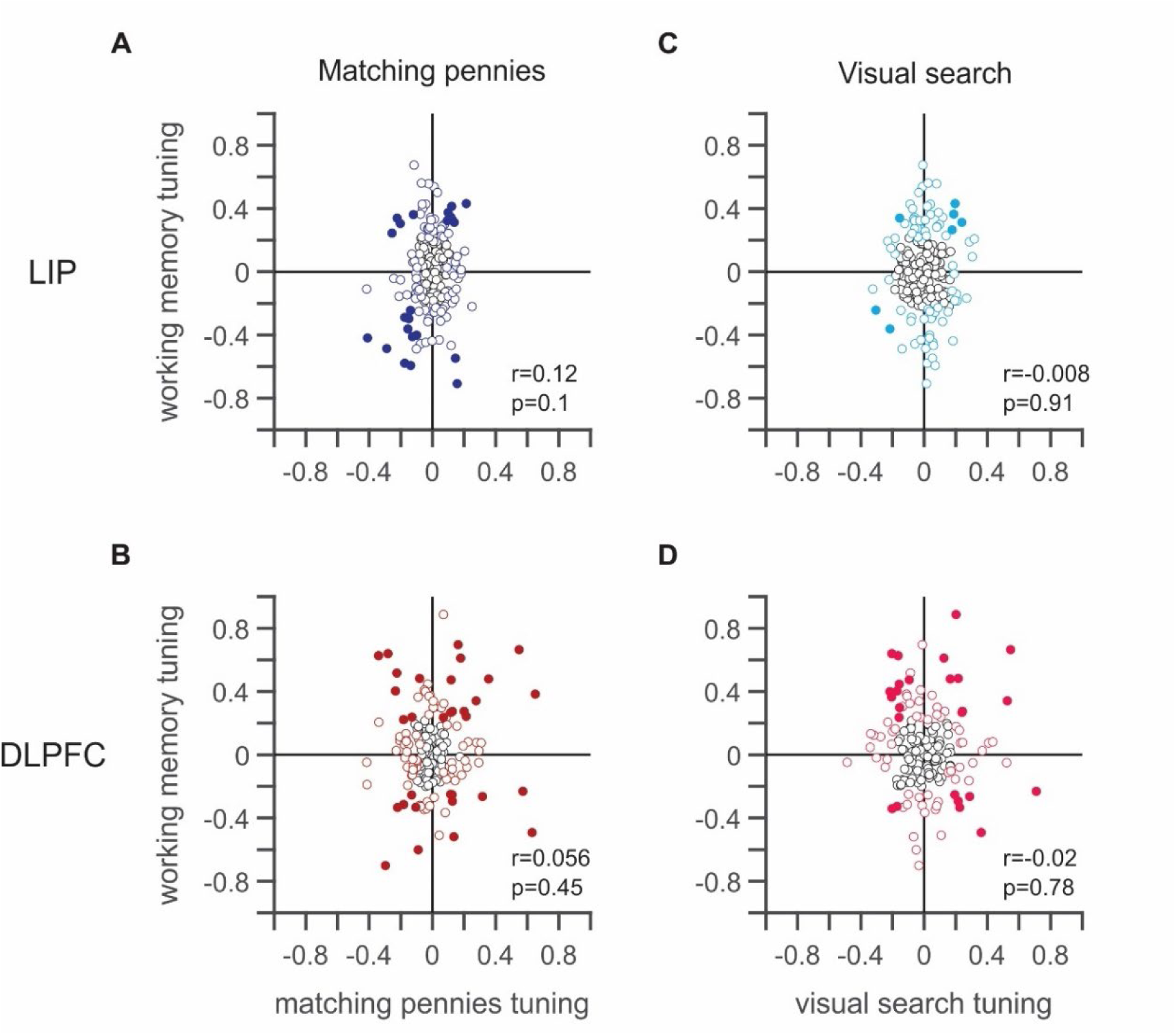
Relationship between memory tuning for previous choice and working memory horizontal tuning. **A-B.** Relationship between SRCs for working memory horizontal tuning and SRCs for previous choice memory in the matching pennies task for neurons in LIP in (A) and DLPFC in (B). **C-D.** Relationship between SRCs for working memory horizontal tuning and SRCs for previous choice memory in the visual search task for neurons in LIP in (C) and DLPFC in (D). Neurons showing either significant working-memory or previous choice-related activity are outlined in color; neurons showing significant encoding of both variables are filled in color. Neurons not showing significant encoding of either variable are outlined in black.

**Figure 4.**
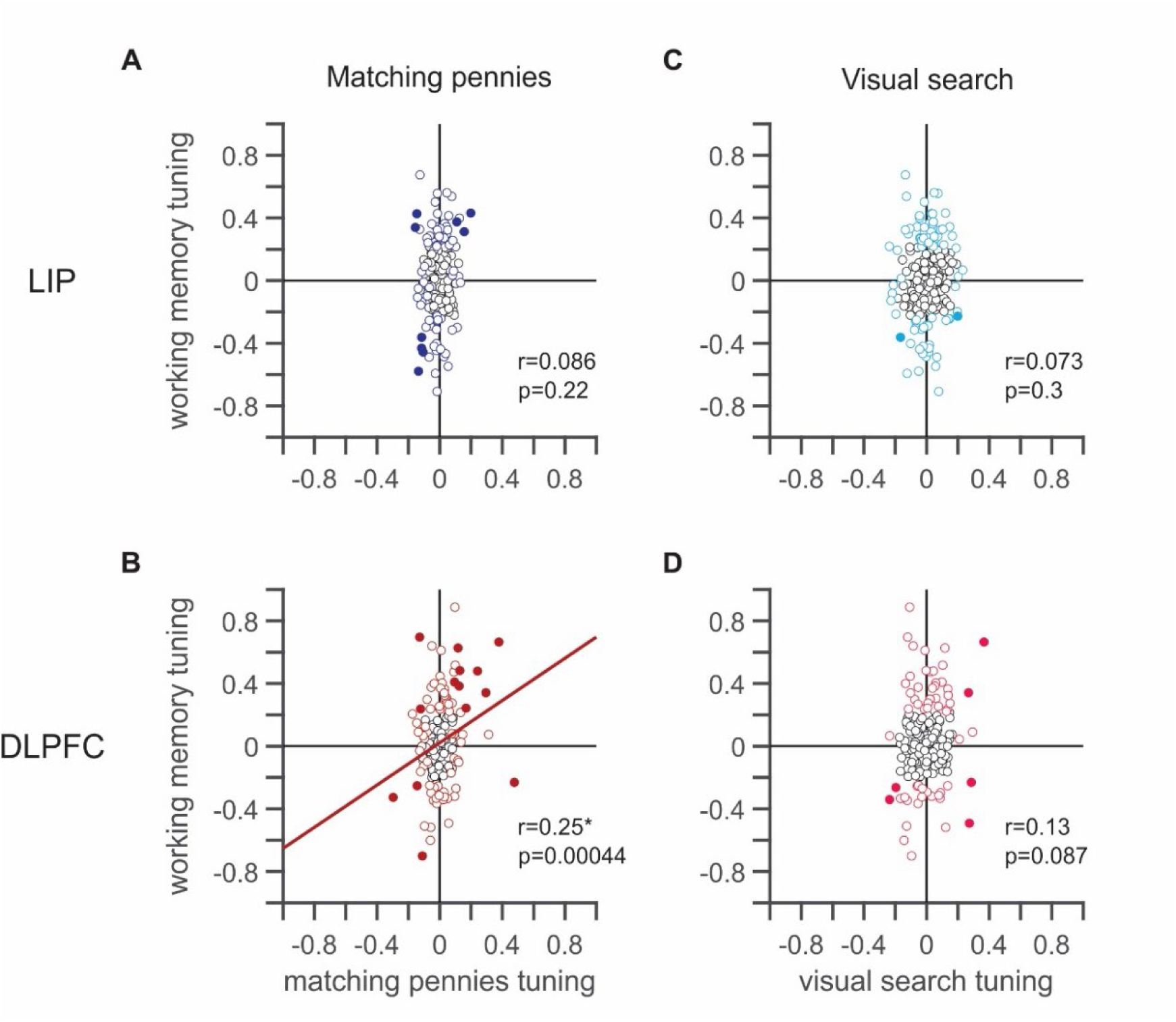
Relationship between memory tuning for previously rewarded choice and working memory horizontal tuning. **A-B.** Relationship between SRCs for working memory horizontal tuning and SRCs for previous choice memory in the matching pennies task for neurons in LIP in (A) and DLPFC in (B). **C-D.** Relationship between SRCs for working memory horizontal tuning and SRCs for previous choice memory in the visual search task for neurons in LIP in (C) and DLPFC in (D). Significance testing corrected for multiple comparisons.

An example DLPFC neuron that demonstrates congruent working memory encoding and encoding of the previously remembered choice during the matching pennies task, but not during the visual search task is shown in **Figure 5**. We did not observe a significant relationship between mnemonic working memory tuning and tuning to previously rewarded choice in either decision-making task in LIP (**Fig. 4A**, matching pennies: Pearson’s *r* = 0.086, *p* = 0.22; **Fig. 4C**, visual search: Pearson’s *r* = 0.073; *p* = 0.3; two-tailed).

**Figure 5.**
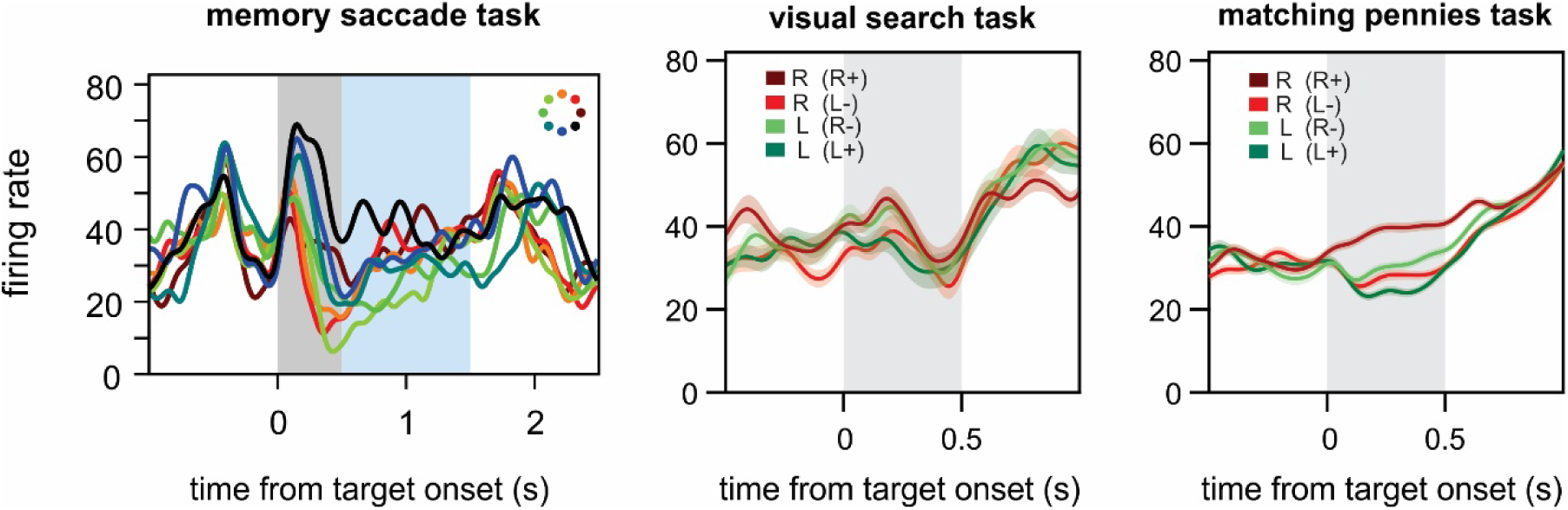
Example DLPFC neuron. Example neuron from DLPFC with activity from all three tasks. The neuron shows congruent (rightward) tuning for the remembered location during the memory saccade delay and for the previously rewarded choice during the delay in the matching pennies task, but not the visual search task.

The correlation between encoding of the previously rewarded choice in matching pennies and working memory tuning was significantly larger in DLPFC compared LIP (Fisher’s *z*-test; *p* = 0.048, one-tailed). The delay period in the decision-making tasks and the memory period in the working memory task both allowed the animal to plan the saccade executed at the end of these epochs. We therefore further hypothesized that single neurons in LIP and DLPFC may encode the upcoming left-right choice decision consistently with mnemonic working memory encoding along the horizontal axis. We confirmed this hypothesis for both decision tasks in LIP, where SRCs for working memory horizontal tuning and the upcoming choice were significantly correlated in both decision-making tasks (**Fig. 6A**; matching pennies: Pearson’s *r* = 0.20, *p* = 0.004; **Fig. 6C**; visual search: *r* = 0.43, *p* = 1.9e-10; two-tailed; corrected for multiple comparisons). This correlation was significantly stronger for the visual search task compared to the matching pennies task (Fisher’s *z*-test; *p* = 0.0049, one-tailed). In DLPFC, we found that there was a significant correlation between working memory tuning and encoding of the upcoming choice for visual search (**Fig. 6D**; Pearson’s *r* = 0.2, *p* = 0.0001; two-tailed; corrected for multiple comparisons). However, this relationship was not present in DLPFC for matching pennies, and the correlation was significantly stronger for visual search compared to matching pennies in DLPFC (**Fig. 6B**; matching pennies: Pearson’s *r* = 0.11, *p* = 0.14, two-tailed; Fisher’s *z* test: *p* = 0.044, one-tailed).

**Figure 6.**
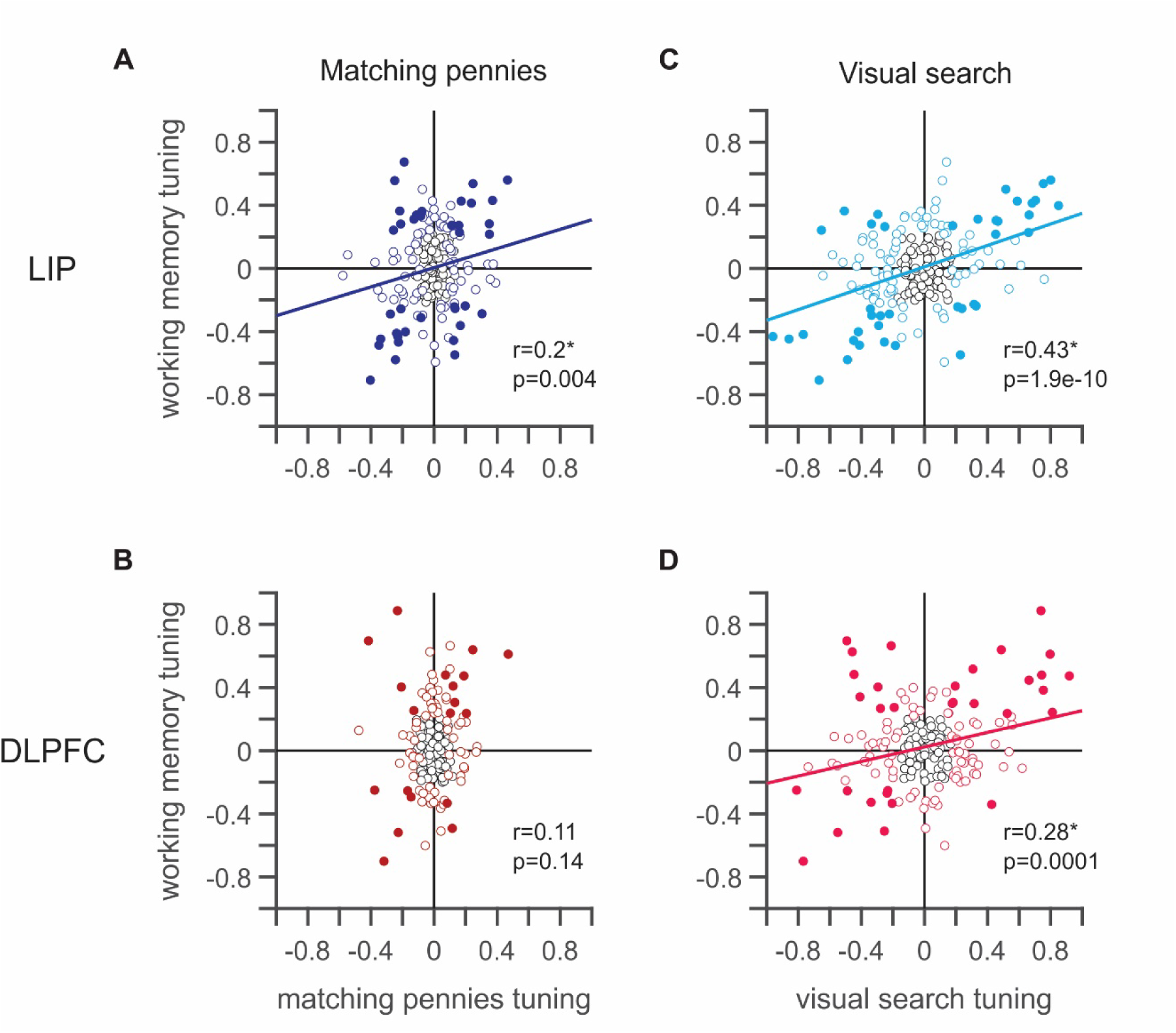
Congruency of encoding of upcoming choice and working memory horizontal tuning. **A-B.** Relationship between SRCs for working memory horizontal tuning and SRCs for encoding of upcoming choice in the matching pennies task for neurons in LIP in (A) and DLPFC in (B). **C-D.** Relationship between SRCs for working memory horizontal tuning and SRCs for encoding of upcoming choice in the visual search task for neurons in LIP in (C) and DLPFC in (D). Significance testing corrected for multiple comparisons.

Given that we did not see consistent encoding of the upcoming saccades across all tasks and regions, we wondered whether seemingly intrinsic neuronal properties might also differ across tasks. We therefore then asked whether the visuomotor response properties of each neuron estimated from the working memory task were consistent with the visuomotor response properties of the same neurons estimated during decision-making. We quantified the visual-motor index (VMI; Ray et al., 2009) for each neuron and task and examined the relationship between these indices for the ODR task and each of the two decision-making tasks.

Surprisingly, we found that in DLPFC, the neurons’ VMIs were consistent and significantly correlated between the ODR task and the visual search task but not between the ODR task and the matching pennies task (**Fig. 7B**; visual search: *r* = 0.45, *p* = 5.0e-11*; **Fig. 7D**; matching pennies: *r* = 0.060, *p* = 0.41; two-tailed; corrected for multiple comparisons). The strength of the correlation between VMIs for visual search and memory saccade was significantly larger than the correlation between matching pennies and memory saccade in DLPFC (Fisher’s *z* test; *p* = 1.1e-5, one-tailed). Furthermore, in LIP, VMIs were not correlated between the ODR task and either of the decision-making tasks (**Fig. 7A**; visual search: r = 0.056, p = 0.42; **Fig. 7C**; matching pennies: *r* = 0.13, *p* = 0.061; two-tailed). The strength of the correlation of VMIs between visual search and memory saccade in DLPFC was also significantly larger than the same correlation in LIP (Fisher’s *z*-test; *p* = 9.6e-6, one-tailed).

**Figure 7.**
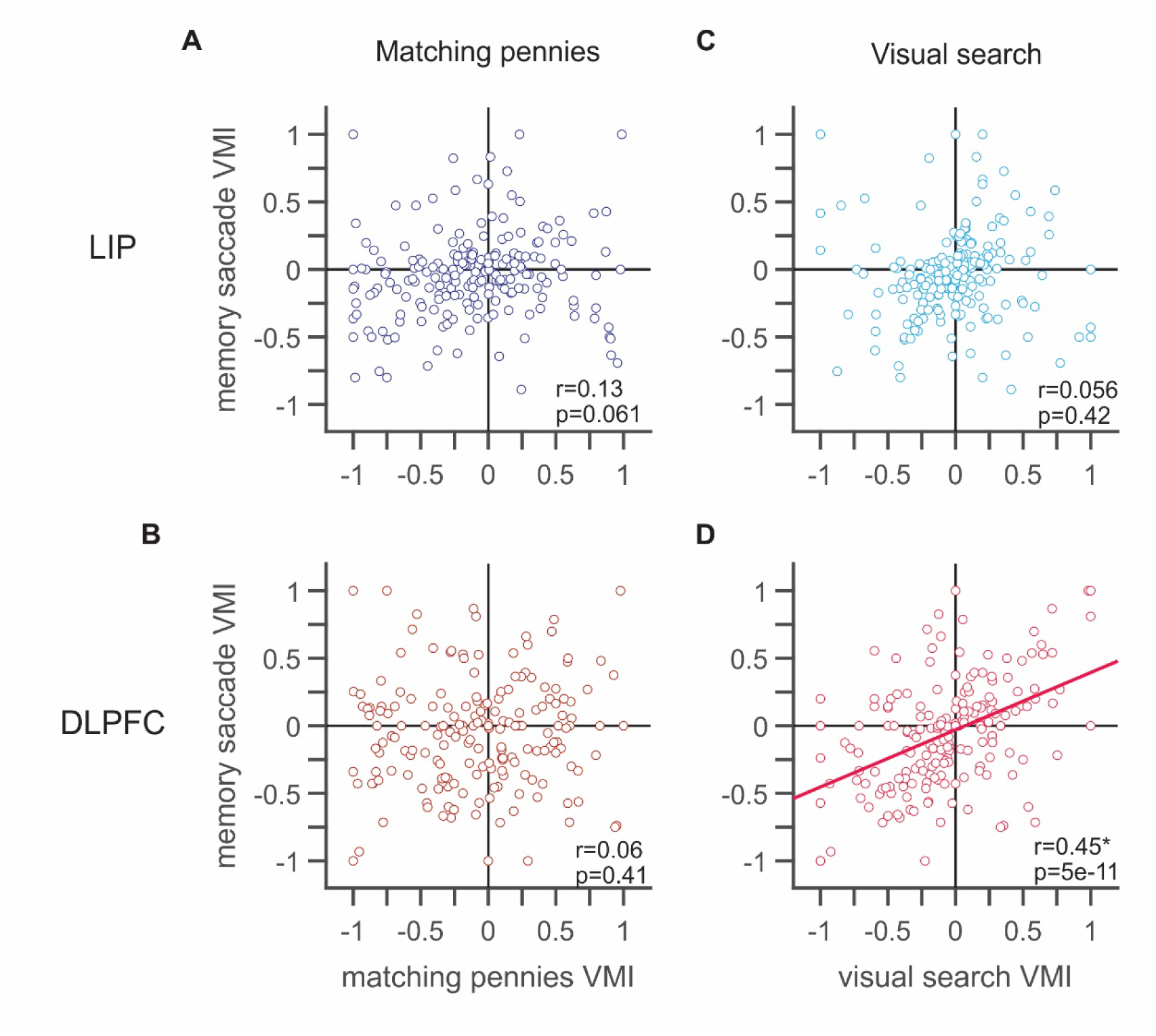
Consistency of visual-motor index between working memory and decision-making. **A-B.** Relationship between VMIs calculated in the ODR task and VMIs calculated in the matching pennies task for neurons in LIP in (A) and DLPFC in (B). **C-D.** Relationship between VMIs calculated in the ODR task and VMIs calculated in the visual search task for neurons in LIP in (C) and DLPFC in (D). Significance testing corrected for multiple comparisons.

## Discussion

In the present study, we examined the consistency between encoding of mnemonic representations in working memory and mnemonic representations for decision-making variables in DLPFC and LIP. A significant fraction of neurons in each region encoded the upcoming and previous choice during the decision period in both decision-making tasks. We also observed a significant fraction of neurons in both regions encoding the previously rewarded choice during the delay period in matching pennies, where the animals used a reinforcement learning strategy, but not in visual search, where the animals selected the green target regardless of events in previous trials.

We found evidence of overlap in mnemonic memory representations for working memory and reinforcement learning in DLPFC, where the neural population tended to congruently encode information in working memory and the mnemonic representation for the previously rewarded choice. This relationship was present for the matching pennies task only, where the previously rewarded choice was behaviorally relevant and was not present in the visual search task or in either decision-making task in the LIP.

Although the previous choice was robustly encoded by neurons in both DLPFC and LIP during the decision period of both decision-making tasks, neurons in these regions did not congruently encode the mnemonic representation of the previous choice and the mnemonic working memory representation. On the other hand, we found that the upcoming choice was encoded congruently with the mnemonic working memory representation for both decision-making tasks in LIP as well as in the visual search task for DLPFC. We therefore observed a dissociation in the congruency of encoding of decision-making representations and working memory representations across tasks and regions. The most behaviorally relevant memory for reinforcement learning, the previously rewarded choice, was encoded congruently with the working memory representation for DLPFC. By contrast, the prospective choice was encoded congruently with the working memory representation for reinforcement learning in LIP only and in both regions for visual search. These results suggest that the network mechanism for persistent activity used for working memory might also support the maintenance of the memory of the previously rewarded choice when it is behaviorally relevant in DLPFC but not LIP.

In addition, we found that the visual-motor index for the same neurons was not generally consistent across working memory and decision-making tasks. We observed no relationship between visual-motor indices for the ODR task and matching pennies task in DLPFC or LIP. By contrast, we observed significant congruent relationship between visual-motor indices for the ODR task and the visual search task in DLPFC but not in LIP. This result suggests that the visual-motor index is not a general feature of single neurons but rather a property that varies based on task demands. Overall, these results suggest a specialized role for the DLPFC in supporting behavior which varies based on task demands.

